# CXCL5-mediated recruitment of neutrophils into the peritoneal cavity of *Gdf15*-deficient mice protects against abdominal sepsis

**DOI:** 10.1101/613901

**Authors:** Isa Santos, Henrique G. Colaço, Ana Neves-Costa, Elsa Seixas, Tiago R. Velho, Dora Pedroso, André Barros, Rui Martins, Nuno Carvalho, Didier Payen, Sebastian Weis, Hyon-Seung Yi, Minho Shong, Luís Ferreira Moita

**Author notes:** Corresponding author: Luis F. Moita. These authors contributed equally to this work (Isa Santos, Henrique G. Colaço, Ana Neves-Costa).

## Abstract

Sepsis is a life-threatening organ dysfunction condition caused by a dysregulated host response to an infection. Here we report that the circulating levels of growth-differentiation factor-15 (GDF15) are strongly increased in septic shock patients and correlate with mortality. In mice, we find that peptidoglycan is a potent ligand that signals through the TLR2-Myd88 axis for the secretion of GDF15 and that *Gdf15*-deficient animals are protected against abdominal sepsis due to increased chemokine CXC ligand 5 (CXCL5)-mediated recruitment of neutrophils into the peritoneum leading to better local bacterial control. Our results identify GDF15 as a potential target to improve sepsis treatment. Its inhibition should increase neutrophil recruitment to the site of infection and consequently lead to better pathogen control and clearance.

## Introduction

Sepsis is a complex disorder caused by a non-adaptive host response to an infection, leading to acute organ dysfunction and consequent high risk of death (1). It is the leading cause of death in intensive care units and the third cause of overall hospital mortality (2). The pathophysiology and molecular basis of sepsis remain poorly understood. The urgently needed novel therapies for sepsis can only be inspired by new insights into the molecular basis of multiple organ failure and endogenous tissue protective mechanisms. There are two evolutionarily conserved defense strategies against infection that can limit host disease severity. One relies on reducing pathogen load, i.e. resistance to infection that courses with and requires inflammation, while the other provides host tissue damage control, limiting disease severity irrespectively of pathogen load, i.e. tolerance to infection (3).

Growth and Differentiation Factor 15 (GDF15), also known as NSAID-activated gene 1 (NAG-1) and macrophage inhibitory cytokine 1 (MIC-1), is a stress response cytokine and a distant member of the transforming growth factor beta (TGFβ) superfamily (4). The role and effects of GDF15 are best documented in obesity and regulation of energy homeostasis (5). Four independent laboratories have recently identified glial-derived neurotrophic factor (GDNF) receptor alpha-like (GFRAL) as a central receptor for GDF15, likely using the tyrosine kinase receptor Ret as a co-receptor (6–9). Signaling through GFRAL accounts for the central actions of GDF15, including suppression of food intake, but it is unlikely to also explain the peripheral effects of GDF15 (10). GDF15 serum levels have been shown to be elevated in many disease processes that course with cellular stress, prominently cancer and cardiovascular disease (11).

Anorexia and weight loss are key features of the wasting syndrome that often accompanies late-stages of cancer and where GDF15 is known to play a critical role (12). These signs, including long-lasting loss of muscle mass, are also observed in sepsis (13). The common pathological features of both conditions led us to investigate whether GDF15 is increased in sepsis and if it plays a role in its pathophysiology.

## Results and discussion

To begin addressing these questions, we measured the level of GDF15 in 40 serum samples from septic patients and compared the results with those of 130 healthy controls (HC) and 33 patients with a histological diagnosis of acute appendicitis (AA, 20 diagnosed with acute phlegmonous appendicitis and 13 with acute gangrenous appendicitis) before surgery. We found significant differences in GDF15 levels both when comparing AA with HC (p<0.0001) and AA with sepsis patients (p<0.0001, Figure 1A). Septic patients had substantially higher values than the other groups (Figure 1A), in good agreement with a previous recent report (14). We then compared survival of septic patients 28 days after diagnosis and found that survivors had on average substantially lower levels of GDF15 on the day of diagnosis (Figure 1B). Importantly, patients with less than 10ng/mL GDF15 at the day of diagnosis showed a low mortality rate (5.8%) 28 days post-diagnosis, whereas those with higher GDF15 levels had a high mortality rate (39.1%, Figure 1C). None of the clinical or analytical severity parameters of AA patients correlated with GDF15 levels (data not shown). When divided based on mortality, sepsis patients did not differ in terms of age (W=191, p=0.2054) or gender (p=1) (Supplemental Table I). Survivors had a lower SAPS II (W=244.5, p=0.0033) and APACHE II score (W=108, p=0.0174), but did not significantly differ on the SOFA score at day 1 (W=188, p=0.0702) as compared to non-survivors.

**Figure 1.**
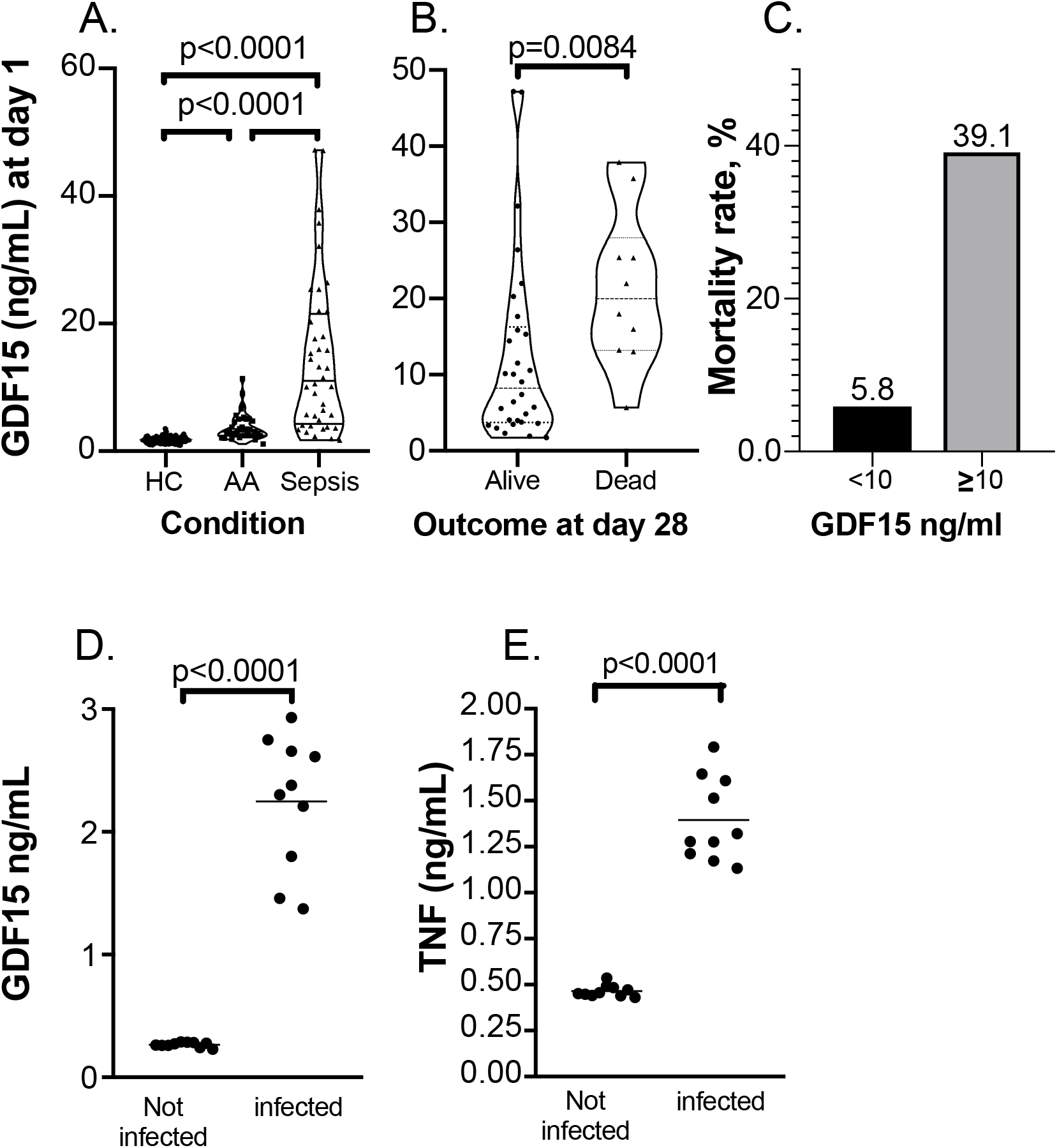
GDF15 is induced by infection in humans and mice. **(A)** Serum levels of GDF15 in acute appendicitis (AA) and severe sepsis compared with healthy controls (HC). **(B)** Serum levels of GDF15 in septic patients at day 1 of admission to intensive care unit (ICU) comparing survival outcome at 28 days after admission to ICU. **(C)** Comparison of mortality rates in patients with GDF15 serum levels lower than 10ng/mL and higher than 10ng/mL at day 1 of admission to ICU. **(D)** and **(E)** GDF15 and TNF serum levels, respectively, in mice infected with *E. coli i.p*. (n=10) or injected with vehicle (n=10), quantified at the peak of cytokine detection.

To explore the role of GDF15 in sepsis, we first measured its levels in the peripheral blood of mice challenged with an i.p. injection of *E. coli* and compared them with vehicle-treated mice 24 h after injections (Figure 1D). We found that similarly to TNF in serum at 2 h after challenge (Figure 1E), the levels of GDF15 were significantly increased (Figure 1D), in agreement with the findings in sepsis patients (Figure 1A). We then used a panel of microbial pattern recognition agonists to challenge bone marrow derived macrophages (BMDM) from wild-type (WT) C56BL/6 mice and found that TLR2 agonists were the strongest inducers of GDF15 expression and secretion (Figure 2A) in contrast with the broad pattern of TNF secretion measured in the same conditioned media (Figure 2A). Based on these results, we chose peptidoglycan (PGN) from *Bacillus subtilis* for further experiments (Figure 2B). In contrast with either *E. coli*, or growing concentrations of lipopolysaccharide (LPS) that robustly induce the secretion of TNF (Supplemental Figure 1A) but do not induce the secretion of GDF15, PGN from *B. subtilis* strongly induces the secretion of GDF15 in a concentration dependent manner (Figure 2B). We then asked which pathway senses PGN leading to the secretion of GDF15. In comparison with BMDM from WT C56BL/6, we found that the TLR2-MyD88 pathway is required for GDF15 secretion while both NOD1 and NOD2 are dispensable (Figure 2C). Depending on the dose and duration of challenge with PGN, TLR4/MyD88 is also capable of sensing PGN to induce GDF15 secretion (Figure 2C and data not shown).

**Figure 2.**
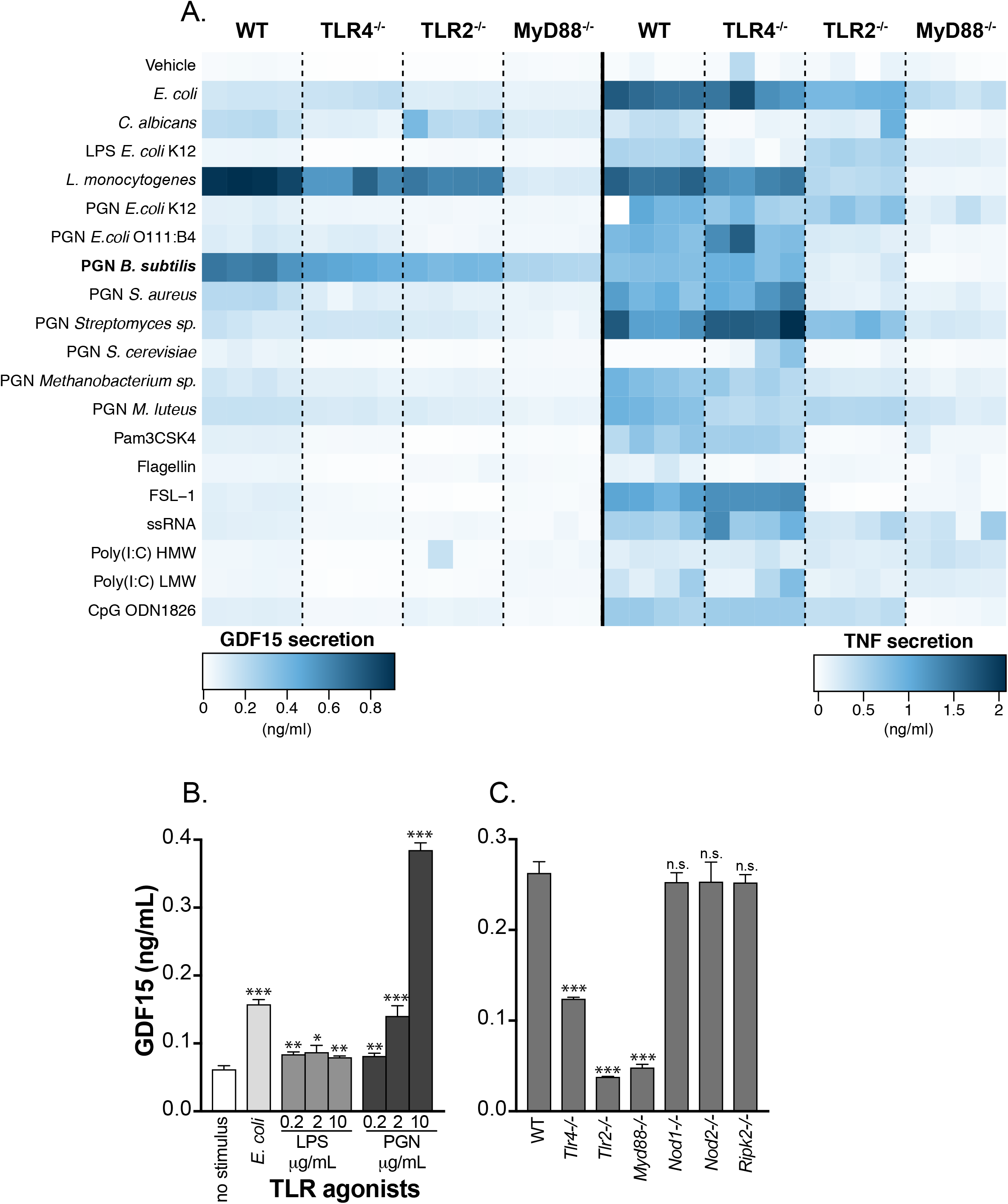
GDF15 is induced by PGN through a TLR2-Myd88 pathway in mice. Quantification by ELISA of GDF15 and TNF secretion induced by TLR agonists in BMDMs. **(A)** GDF15 and TNF levels in conditioned media following incubation of BMDMs from wild-type (WT) or knockout animals (*Tlr4^−/−^, Tlr2^−/−^* and *Myd88^−/−^*) for 24 h without stimulus or with a series of TLR agonists. TLR agonists and concentrations: PFA-fixed *E. coli* at 20 bacteria per cell; PFA-fixed *C. albicans* at 20 yeast per cell; LPS from *Escherichia coli* at 200ng/mL; *Listeria monocytogenes* at 100 million bacteria/mL; peptidoglycan (PGN) from a variety of microbial sources at 10, 5 and 2.5μg/mL; Pam3CSK4 at 300ng/mL; flagellin from *S. typhimurium* at 1μg/mL; FSL-1 (Pam2CGDPKHPKSF) at 100ng/mL; ssRNA40 at 2.5μg/mL; Poly(I:C) HMW at 10μg/mL; Poly(I:C) LMW at 10μg/mL; and CpG oligonucleotide (ODN) at 1.5μM. **(B)** GDF15 levels in conditioned media following incubation of WT BMDMs for 24 h without stimulus or with the TLR agonists: PFA-fixed *E. coli* at a ratio of 20 bacteria per cell; lipopolysaccharide (LPS) from *E. coli* at 0.2 to 10μg/mL; and peptidoglycan (PGN) from *B. subtilis* at 0.2 to 10μg/mL. **(C)** GDF15 levels quantified as in B, following incubation of BMDMs from WT or knockout animals (*Myd88^−/−^*, *Tlr2^−/−^*, *Tlr4^−/−^*, *Nod1^−/−^*, *Nod2^−/−^* and *Ripk2^−/−^*), for 16 h with PGN 2.5μg/mL. Averages and standard deviations are shown for three replicate wells in 96-well plates; one representative experiment is shown and at least three independent experiments were performed and quantified by ELISA.

We then turned to the mechanistic role of GDF15 in sepsis as the strongly increased secretion could either mean this factor is involved in a compensatory response to a stress (infection) challenge or that it plays an active part in increasing the severity of infection. To distinguish between these two possibilities, we compared the survival rates of WT and *Gdf15*-deficient (*Gdf15^−/−^*) mice in response to a polymicrobial peritonitis using the cecum ligation and puncture (CLP) model of sepsis. In multiple and independent experiments, we consistently observed that *Gdf15^−/−^* animals are protected against sepsis and survive for considerably longer periods (Figure 3A). Sickness behavior is defined as a group of signs and symptoms, including anorexia, lethargy, fever, altered sleep patterns, lack of grooming, and social withdrawal in response to an infection. In line with this, *Gdf15^−/−^* mice showed reduced temperature loss as compared to WT mice, both 8h and 24h post-CLP (Figure 3B). *Gdf15^−/−^* mice have lower scores for sickness behavior, as detailed in Supplemental Figure 2. All tested cytokines had lower levels in *Gdf15^−/−^* animals (Supplemental Figure 3), which reached statistical significance in the cases of Interleukin (IL)-1β and IL-12 (Supplemental Figure 3C and F) suggesting a better control of infection in *Gdf15^−/−^* animals. We found no relevant differences in the levels of serologic markers of organ dysfunction or damage (including creatinine, LDH, CK, AST and ALT) at 24 h, suggesting organ damage was similar in both strains (Supplemental Figure 4A, B, C, D and E). This result is in agreement with minor histological differences between both genotypes at 24 h after CLP (Supplemental Figure 4F).

**Figure 3.**
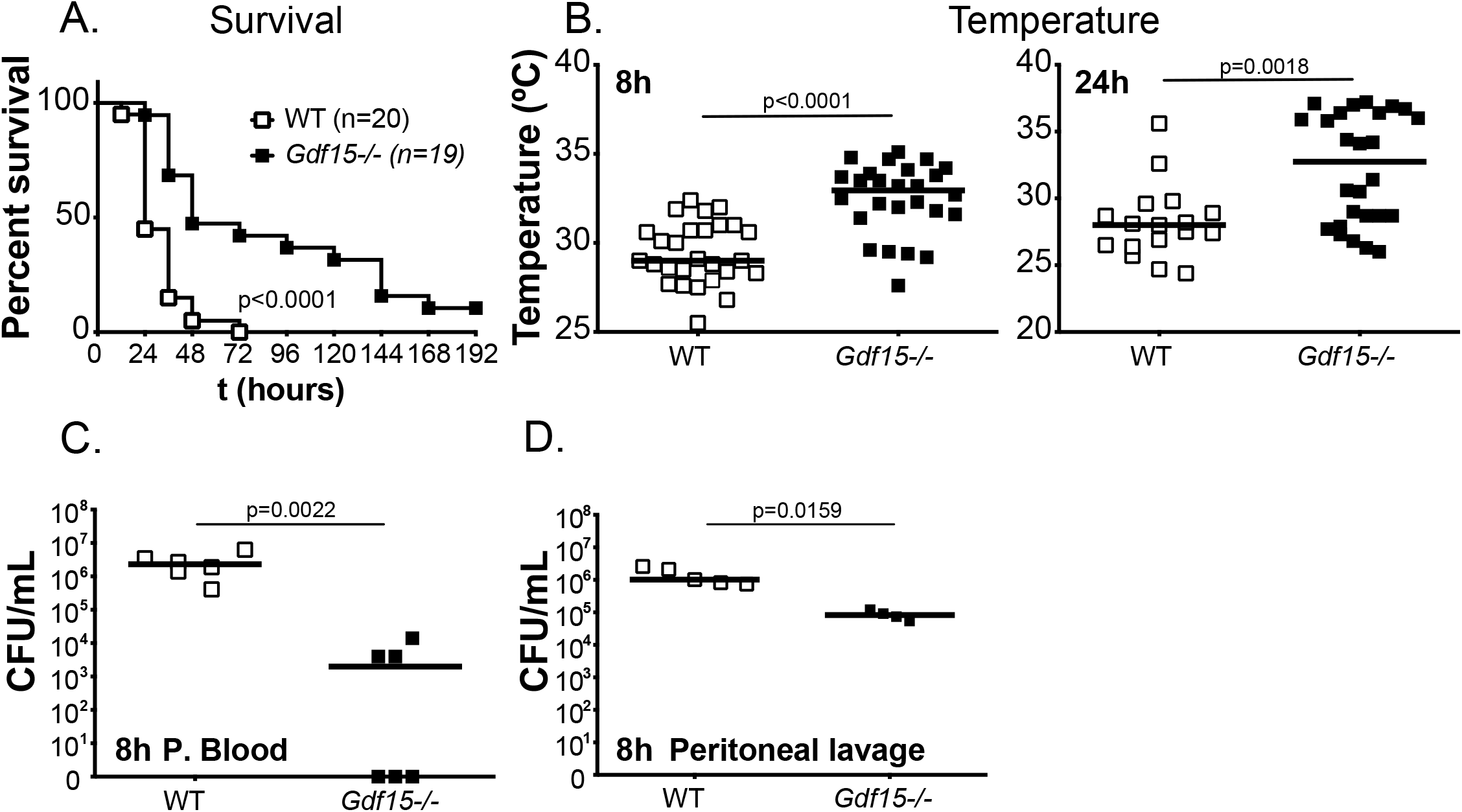
*Gdf15^−/−^* mice are protected against CLP and have decreased CFUs in the peritoneum. **(A)** Survival to cecal ligation and puncture (CLP) comparing wild-type (WT, n=20) and *Gdf15^−/−^* animals (n=19). **(B)** Rectal temperature in WT (n=28) and *Gdf15^−/−^* (n=26) animals 8 and 24 h after CLP. **(C)** CFUs cultured from peripheral blood of WT (n=6) and *Gdf15^−/−^* (n=6) animals 8 h after CLP. **(D)** Colony-forming units (CFUs) cultured from peritoneal lavage of WT (n=5) and *Gdf15^−/−^* (n=4) mice 8 h after CLP.

We then measured the bacterial load in the peripheral blood of WT and *Gdf15^−/−^* animals in independent experiments and found consistently a statistically significant lower bacterial burden in *Gdf15^−/−^* mice at 8 h after CLP (Figure 3C), suggesting that *Gdf15^−/−^* better control the initial local infection. Together, the results for cytokines, serologic markers of organ lesion, histopathology and bacterial burden, suggest that *Gdf15^−/−^* mice are more resistant to infection, without affecting disease tolerance, as defined in (15). To investigate further this possibility, we analyzed and compared the peritoneal lavage contents of WT and *Gdf15^−/−^* mice after 8 h of CLP. We found bacterial levels on the peritoneal lavage to be on average 10-fold lower (p=0.0159) in *Gdf15^−/−^* mice (Figure 3D), which was in good agreement with substantially elevated relative (Figure 4A, p<0.05) and absolute (Figure 4B, p<0.05) numbers of neutrophils. The increased number of neutrophils, rather than their differential activity between genotypes, is likely to be responsible for a better local control of the initial infection as we did not observe increased phagocytic activity (data not shown). These findings are consistent with previous observations in a mouse model of myocardial infarction where *Gdf15^−/−^* mice recruited more neutrophils to the injury site (16). In this model, mice had a worse outcome as the excess number of neutrophils exacerbated the myocardial lesion (16). An excess recruitment of neutrophils has also been observed in *Gdf15^−/−^* mice in a model of CCl_4_-induced liver fibrogenesis (17), which contributed to lesion exacerbation. In contrast, in the present case of the CLP model the increased number of neutrophils lead to the better early control of the local infection. Taken together, our findings and previously published observations, converge on the regulation of neutrophil recruitment, supporting a role for GDF15 in the regulation of tissue injury and that its effects are likely to be highly contextual.

**Figure 4.**
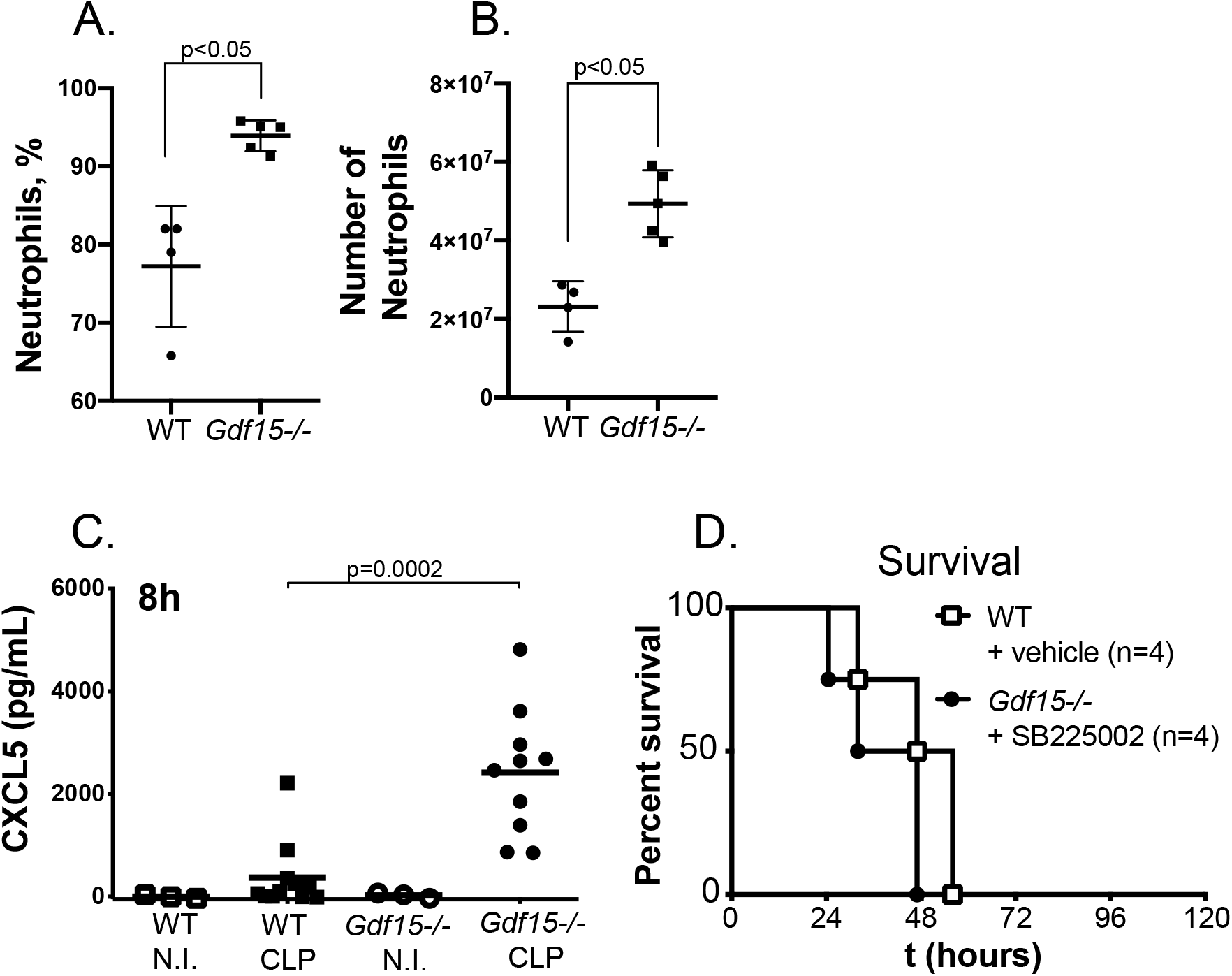
*Gdf15^−/−^* mice better control local infection due to CXCL5-mediated neutrophil influx into the peritoneum. **(A)** Percentage and **(B)** total number of neutrophils in the peritoneal lavage of WT (n=4) and *Gdf15^−/−^* (n=5) mice 8 h after CLP. **(C)** Quantification by ELISA of CXCL5 in the peritoneal lavage 8 h after CLP in WT and *Gdf15^−/−^* mice without (non-infected, NI) or 8 h after CLP. **(D)** Survival to CLP comparing WT vehicle-treated (WT, n=4) and *Gdf15^−/−^* mice treated with CXCR2 inhibitor SB225002 (*Gdf15^−/−^*, n=4).

To identify mechanistic bases for the different numbers of neutrophils at the site of infection, we measured the main chemokine factors for neutrophil attraction, including the CCR1 ligands CCL3 (Supplemental Figure 5 Panel A), CCL4 (Panel B) and CCL5 (Panel C) and the CXCR2 ligands MIP-2/CXCL-2 (Supplemental Figure 5 Panel D), KC/CXCL-1 (Panel E) and CXCL5 (Figure 4C). GDF15 was increased systemically only in WT animals and completely absent in knockout animals (Supplemental Figure 5 Panel F). IL-6 and MCP-1, a chemokine for macrophages, were very similar suggesting that CLP procedure was made with similar severity in both genotypes (Supplemental Figure 5 Panels G and H respectively).

Only LIX/CXCL5 showed a strong, consistent and significant increase in *Gdf15^−/−^* mice (p=0.0002), while all the other chemokines were either very similar or even marginally decreased. To test whether increased signaling through the CXCL5-CXCR2 axis is directedly responsible for the *Gdf15^−/−^* protective phenotype, we used a CXCR2-selective antagonist, SB225002, and compared survival of WT and *Gdf15^−/−^* mice after CLP and treatment with this CXCR2 antagonist. In these conditions, the survival advantage of *Gdf15^−/−^* mice to CLP is lost (Figure 4D), suggesting that increased local levels of CXCL5 in *Gdf15^−/−^* mice are a necessary component of their increased survival to a peritoneal polymicrobial infection.

The significance of our findings is highlighted by the knowledge that neutrophils are critical innate immune cells that provide the first line of host defense against sepsis through their ability to rapidly migrate to the site of infection; in addition, many animal and clinical sepsis studies have indicated that neutrophils display markedly impaired recruitment to infectious sites and fail to clear bacteria, resulting in excessive inflammation and increased mortality (18), findings that lacked mechanistic bases before our findings. Our results raise the possibility that GDF15 may play a key detrimental effect in sepsis by inhibiting neutrophil recruitment to the site of infection, which constitutes an important immunosuppression component that characterizes late stages of sepsis and is responsible for the inability to control and clear infection (19). Strategies to reverse the defects in migration of neutrophils have been proposed as promising therapies for sepsis (20). Our work indicates GDF15 as a possible target for this goal.

Another important implication of our work relates to the identification of CXCL5 as the causal mediator responsible for increased survival of *Gdf15^−/−^* mice, suggesting that the expression of this chemokine is suppressed by GDF15. CXCL5 has been directly implicated in obesity and insulin resistance (21). In the context of our findings, this fact is particularly interesting because it might mean that the obesity phenotype of *Gdf15-/-* mice does not depend exclusively on absence of signaling through the central GFRAL central receptor, but also on a peripheral inflammatory component mediated by CXCL5. In fact, while *Gdf15-/-* animals have increased body weight when fed a standard chow diet, mice lacking GFRAL do not differ in body weight or composition when compared with WT animals fed a standard diet (7–9).

In conclusion, our results suggest that *Gdf15^−/−^* mice are better able to control the initial infection by locally secreting higher levels of the CXCL5 chemokine, known to be involved in neutrophil recruitment through the activation of the CXCR2 receptor (22).

## Author contributions

IS and HGC performed all mouse experiments with the help of ES, AB and TV. ANC and DP performed all experiments in BMDM. HGC, ANC, DP, NC, SW and DP measured biochemical parameters in patients and mouse serum, mouse peritoneal lavage, and BMDM supernatants. ES analyzed mouse peritoneal lavage cells by flow cytometry. RM performed phagocytic assays in BMDM. AB did the statistical analysis of the data. HSY and MS provided reagents and advised the project. ANC contributed to the supervision of the study. LFM conceived the study, coordinated all experiments and wrote the paper with input from all authors.

## Acknowledgements

We are grateful to Gabriel Núñez (University of Michigan, MI 48109, USA), Ramón Merino Pérez (IBBTEC, 39011 Santander, Spain) and Maria Fernández Velasco (CSIC-UAM, 28029 Madrid, Spain) for the kind gifts of *Nod1*-*Nod2*- and *Ripk2*-deficient mice. HGC is supported by a fellowship from Fundação para a Ciência e Tecnologia (FCT, SFRH/BD/105998/2014). L.F.M. is an FCT Investigator and is supported by the European Community Horizon 2020 (ERC-2014-CoG 647888-iPROTECTION) and FCT (FCT: PTDC/BIM-MEC/4665/2014).

## Methods

Further information can be found in the Supplemental Methods and in Supplemental Figures 1–5.

### Statistical analysis

Mann-Whitney test and Mantel-Cox test (for survival curves) were used for statistical analysis, using Graphpad Prism 6.0 (GraphPad Software). A p-value <0.05 was considered statistically significant.

### Study approval

#### Human studies

Human sepsis samples were included under a project approved by The French Ethical committee (‘‘Comité de Protection des Personnes’’ Ile de France IV; 2010-A0004039). Samples from healthy persons and patients with acute appendicitis patients were included under a research project approved by the Ethics Committee of Garcia de Orta Hospital, Portugal (Reference 05/2015). Written informed consent was obtained from either each enrolled patient or their families. The work has been carried out in accordance with The Code of Ethics of the World Medical Association (Declaration of Helsinki). The authors declare no conflict of interest.

#### Mice

All animal studies were performed in accordance with Portuguese regulations and approved by the Instituto Gulbenkian de Ciência ethics committee and DGAV (Reference A002.2015).

## Supplemental Figures

**Supplemental Figure 1.**
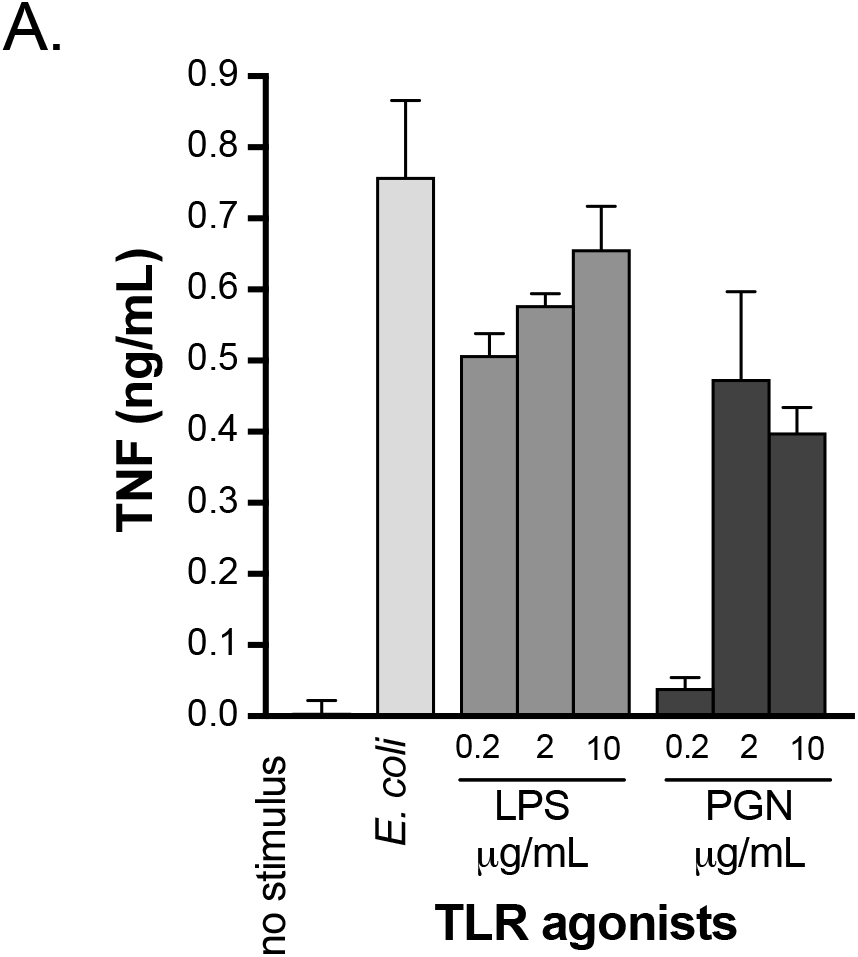
TLR agonists induce GDF15. **(B)** Quantification of TNF levels in the same conditions as in **Figure 2B**.

**Supplemental Figure 2.**
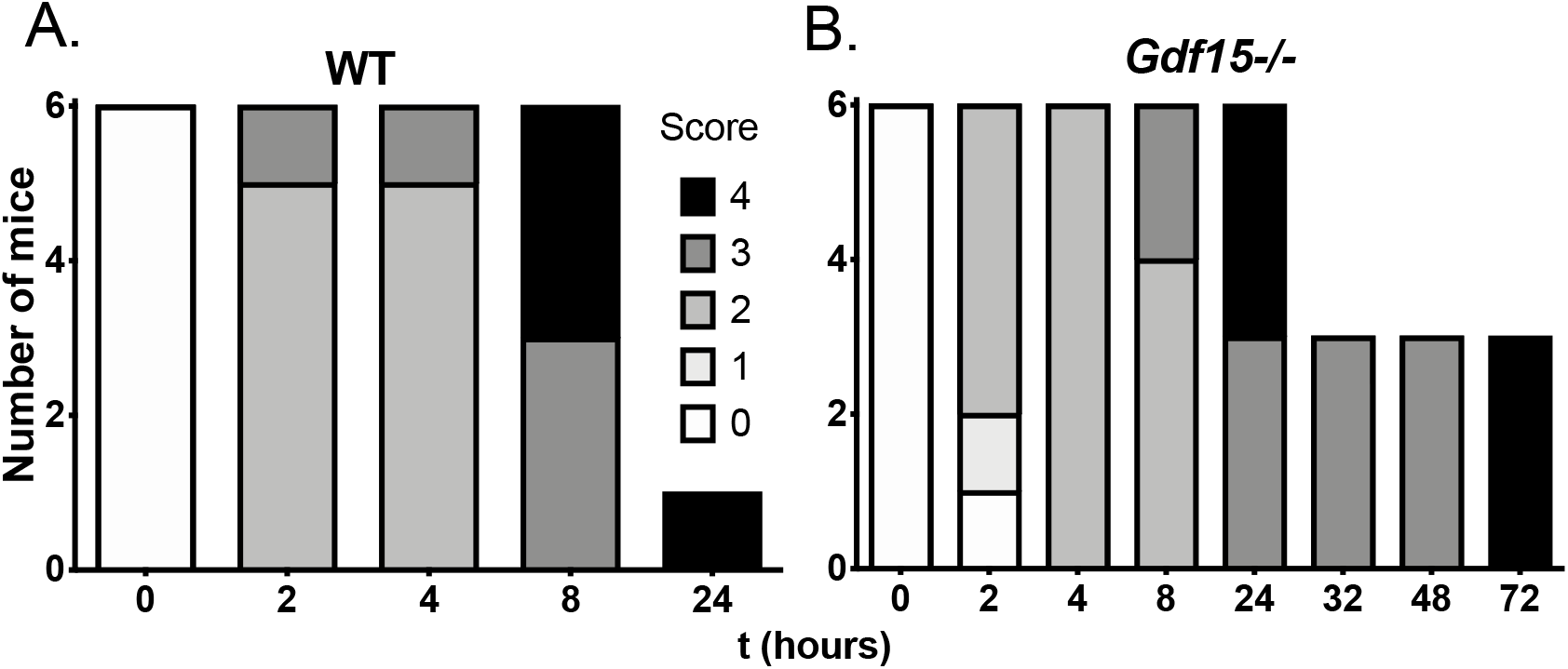
GDF15 modulates sickness behavior. A five-point scale to evaluate the development of sickness behaviors in WT **(A)** and *Gdf15^−/−^* mice **(B)**. At specific time points following CLP, animals were examined by two observers who independently scored the presence of specific signs of sickness. In each animal, the following four signs were evaluated: (1) Piloerection, (2) Ptosis, (3) Lethargy and (4) Huddling. All animals were observed inside their cages. Each parameter was score as presence (1) or absence (0). Graphs represent the distribution of sickness behavior presented at each timepoint. “0” = no sickness behavior; “4” = all four signs of sickness behavior.

**Supplemental Figure 3.**
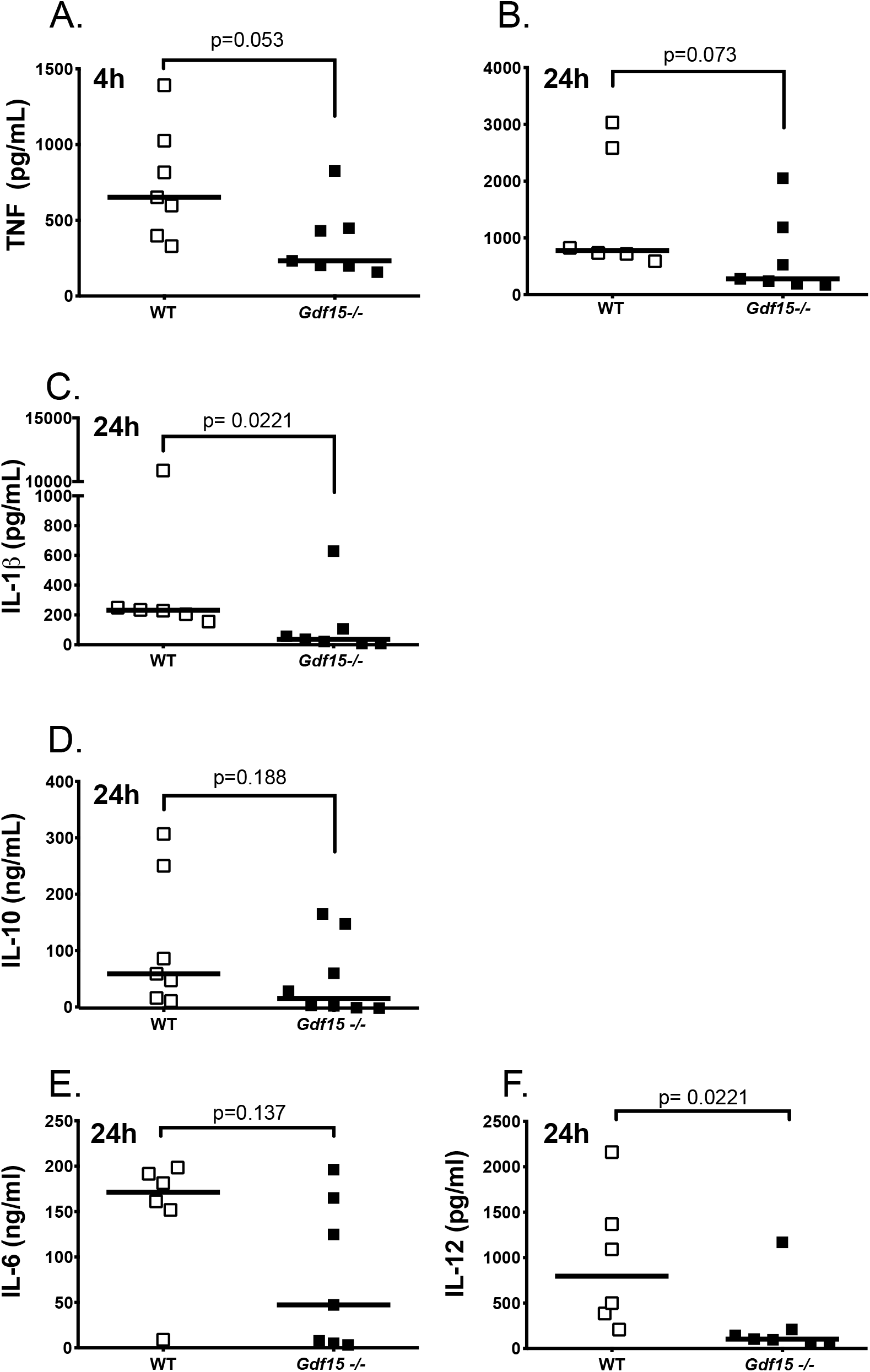
Cytokine quantification in peripheral blood of animals after CLP. **(A)** Quantification by ELISA of TNF at 4 h, **(B)** TNF at 24 h, **(C)** IL-1β at 24 h, **(D)** IL-10 at 24 h, **(E)** IL-6 at 24 h and **(F)** IL-12 at 24 h after CLP in WT or *Gdf15^−/−^* mice.

**Supplemental Figure 4.**
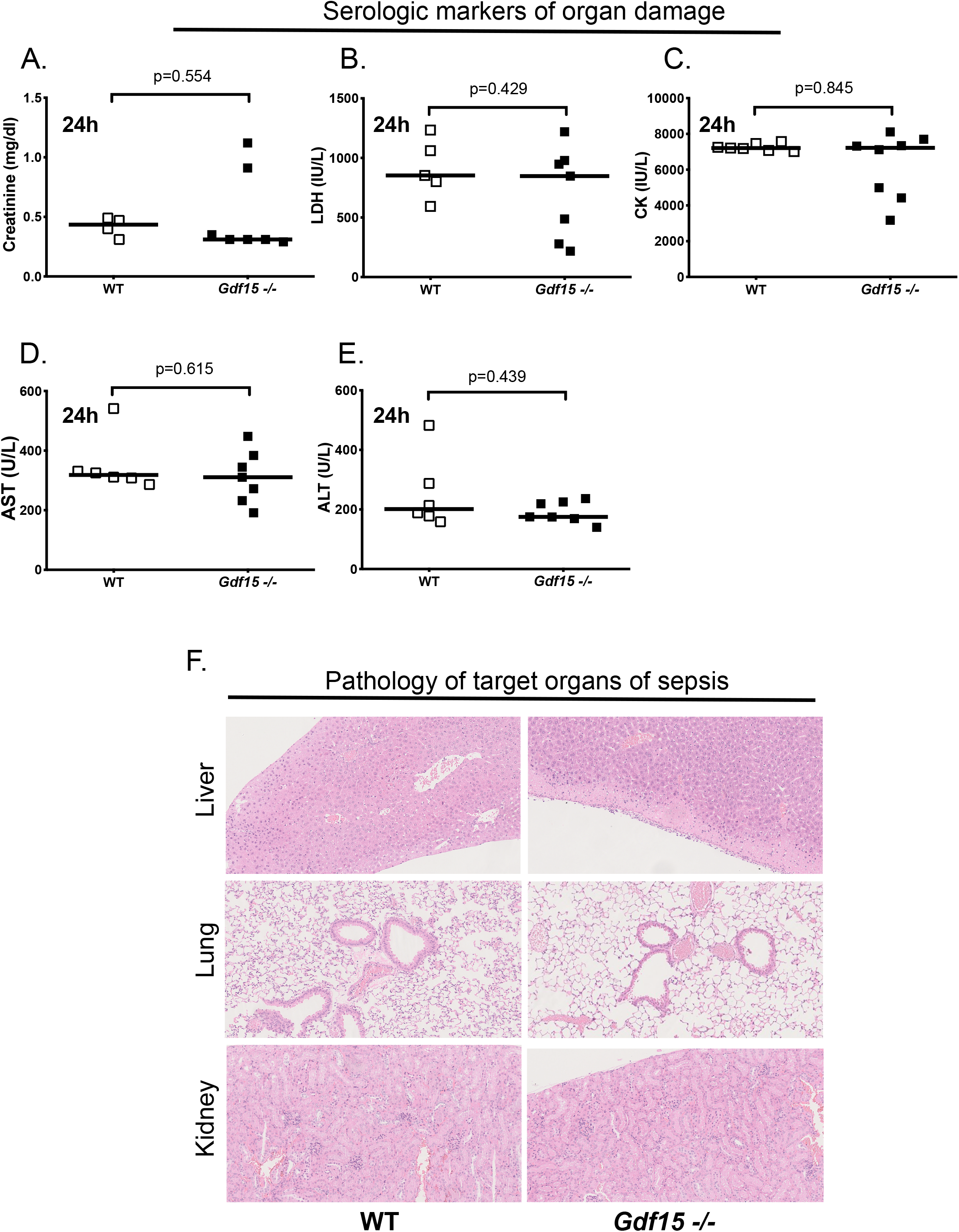
WT and *Gdf15^−/−^* mice have similar degrees of tissue damage after CLP. **(A)**, Colorimetric quantification of serum levels of creatinine, (**B**) LDH, (**C**) CK, (**D**) AST and (**E**) ALT 24 h after CLP in WT and *Gdf15^−/−^* mice. (**F**) Histology analysis of HE stains of liver, lung and kidney from WT and *Gdf15^−/−^* mice 24hr after CLP.

**Supplemental Figure 5.**
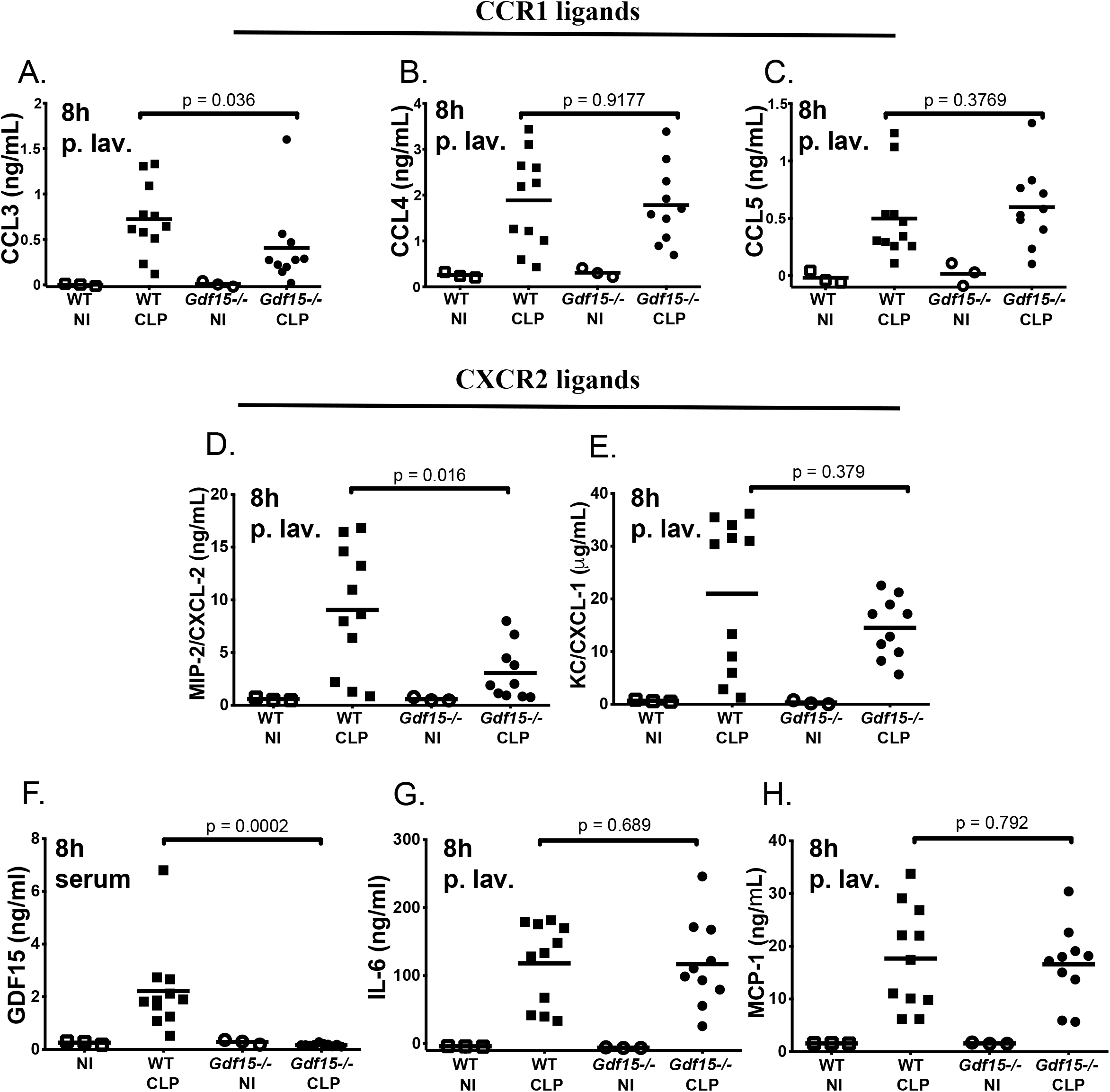
Cytokine and chemokine quantification in the peritoneal lavage and peripheral blood from mice after CLP. Quantification by ELISA of the CCR1 ligands (**A**) CCL3, (**B**) CCL4, (**C**) CCL5 and the CXCR2 ligands (**D**) MIP-2/CXCL-2 and (**E**) KC/CXCL-1 in the peritoneal lavage of WT and *Gdf15^−/−^* mice 8 h after CLP. Quantification by ELISA of (**F**) GDF15, (**G**) IL-6 and (**H**) MCP-1 from WT and *Gdf15^−/−^* mice 8 h after CLP.

## Supplemental Methods

### Mice

*Gdf15* null mice were obtained from M. Shong (Chungnam National University School of Medicine, Daejeon, South Korea). *Gdf15^−/−^*, *Myd88^−/−^*, *Tlr2^−/−^*, *Tlr4^−/−^* and C57BL/6J control mice were bred and maintained under specific-pathogen free conditions at the Instituto Gulbenkian de Ciência with 12 h light/12 h dark cycle, humidity 50–60%, ambient temperature 22 ± 2°C and food and water *ad libitum*.

### Cecal ligation and puncture

Bedding of C57BL/6J and *Gdf15^−/−^* mice was mixed two weeks before the start of the experiments to mitigate differences in microbiome composition. Polymicrobial sepsis was induced in mice by cecal ligation and puncture (CLP), as previously described(23).

### Sickness behavior assay

The Sickness behavior assay used in this report has been described in(24). Briefly, mice were observed by two researchers and rated for the presence (1) or absence (0) of each of the following symptoms: piloerection, ptosis, lethargy, and huddling. Score was presented as the sum of all four parameters.

### *E. coli* challenge in mice

*Escherichia coli* K12 MG1655 was grown in Luria-Bertani broth at 37°C, 180 rpm for 2.5 h until late exponential phase was reached (OD_600nm_=0.8-1.0). The culture was then centrifuged at 4400 xg for 5 minutes at room temperature, washed in the same volume of PBS and centrifuged again. The bacterial pellet was finally resuspended in PBS to obtain an approximately 3×10^9^ CFU/mL. This bacterial suspension was immediately injected intraperitoneally in mice (200 μL/mouse).

### Colony Forming Units assay

Colony forming units (CFU) were determined in blood and peritoneal lavage by serially diluting in sterile PBS and plating in trypticase soy agar plates with 5% sheep blood. Four dilutions were plated per condition. CFU were counted after incubating plates at 37 °C for 16h.

### BMDM treatments

BMDMs were plated in supplemented RPMI 1640 medium (Gibco) and treated 16 h later for the indicated times. The agonists used were from the mouse TLR1-9 Agonist kit (Invivogen tlrl-kit1mw) and peptidoglycan (PGN) was obtained from a variety of sources (all PGN reagents are from Sigma-Aldrich and from a series of microbial origins with the following catalog numbers: from *Bacillus subtilis*, 69554; from *Staphylococcus aureus*, 77140; from *Micrococcus luteus*, 53243; from *Streptomyces* sp., 79682; from *Methanobacterium* sp. 78721; from *Saccharomyces cerevisiae*, 72789). PFA-fixed *Escherichia coli* were incubated with the BMDMs at a ratio of 20 bacteria per cell for the indicated times.

### Biochemical assays in serum, peritoneal lavage, and supernatant from BMDMs

Levels of human GDF15 were quantified in the serum of patients using the ELISA kit DuoSet human GDF15 (R&Systems, catalog number DY957) according to the company’s protocol. Whole mouse blood (obtained by cardiac puncture) was centrifuged at 1600 *xg* for 5 minutes at 4 °C, serum was collected and stored at −80 °C before analysis. Peritoneal lavage was centrifuged at 300 xg for 5 minutes at 4 °C, the supernatant was collected and stored at −80 °C before analysis. For neutrophil identification we used the following antibodies: CD11b, Ly6G, Ly6C, GR1 and anti-neutrophil (7/4). To calculate the absolute number of cells by flow cytometry, we used Perfect-Count Microspheres. The absolute number of the cell population of interest is determined by dividing the number of cells of interest acquired by the number of Perfect-Count Microspheres acquired and multiplying this result by the microsphere concentration. Cytokine and chemokine levels were determined using the following ELISA kits, according to the manufacturer’s instructions: mouse TNF-α (#430902, Biolegend), IL-1β (#432601, Biolegend), IL-6 (#431302, Biolegend), IL-10 (#431411, Biolegend), IL-12/IL-23 (p40) (#431602, Biolegend), CKCL-5/LIX (#DY443, R&D Systems), MCP-1 (#432701, Biolegend), CCL3/MIP-1α (#DY450, R&D Systems), Mouse CCL4/MIP-1β (#DY451, R&D Systems), Mouse CCL5/RANTES (#DY478, R&D Systems), CLCX-1/KC (#DY453, R&D Systems), CXCL-2/MIP-2 (#DY452, R&D Systems), mouse GDF-15 (#DY6385, R&D Systems). Serological makers of organ damage/dysfuntion were determined using the following colorimetric kits, according to the manufacturer’s instructions: QuantiChrom Creatinine (#DICT, Bioassay Systems), QuantiChrom Lactate Dehydrogenase (#D2DH, Bioassay Systems), EnzyChrom Creatine Kinase (#ECPK, Bioassay Systems), EnzyChrom Alanine Transaminase (#EALT, Bioassay Systems), EnzyChrom Aspartate Transaminase (#EALT, Bioassay Systems). All absorbance readings were performed in 96-well plates using an Infinite M200 plate reader (Tecan).

### Flow cytometry

Cells were incubated with different antibodies diluted in PBS containing 2% FBS and 0.01% NaN_3_. To prevent non-specific binding, cells were incubated at 4°C for 20 minutes with CD16/CD32 to block Fc receptors. Afterwards, cells were incubated for 30 minutes at 4°C with the following fluorescent labelled antibodies: anti-CD49b-PE, anti-CD3-PE, anti-B220-PE, anti-Ly6C-APC/Cy7, anti-Ly6G-APC, anti-F4/80-FITC, anti-CD11b-PerCp, anti-CD11b-BV785, anti-F4/80-A647, anti-GR1-PerCp/Cy5.5, anti-Neutrophil (7/4)-FITC, Sytox Blue. All antibodies are from Biolegend except anti-Neutrophil (7/4) which is from Abcam and B220, CD49b and GR1 which are from BD Biosciences. Flow cytometry data were acquired on LSR Fortessa X20 (Becton Dickinson) and analyzed using the FlowJo software package (Tree Star).

### Histopathology

Mouse liver, lung, and kidney were collected 24h after CLP and immediately fixed in 10% buffered formalin. Samples were then embedded in paraffin, sectioned (3 μm) and stained for hematoxilin and eosin according to standard procedures. Histopathology analysis was performed by a trained pathologist, blinded to the mice strain and infection challenge, at the Instituto Gulbenkian de Ciência Histopathology Unit. Tissues were scored for damage, namely necrosis and leukocyte infiltration.

**Supplementary Table I.** Sepsis cohort characterization. Separation based on mortality at day 28.

| Variable | Non-survivors (n=10) | Survivors (n=30) | Statistical Testing |
| --- | --- | --- | --- |
| Age | 75.8 ± 13.39; 77 (72; 85.5) | 65.73 ± 20.79; 71 (58.5; 80) | W = 191<br>p = 0.2054 |
| Gender | 4 Male : 6 Female | 14 Male : 16 Female | Fisher Exact Test;<br>p=1 |
| SAPS II | 64.40 ± 11.18; 65.5 (60.25; 70.50) | 50.6 ± 13.15; 53 (43.50; 58.50) | W = 244.5<br>p = 0.003289 |
| APACHE II | 31.71 ± 9.11 (3 NA); 34 (28; 35.50) | 23.79 ± 4.84 (11 NA); 25 (21.5; 27) | W = 108<br>p = 0.01741 |
| SOFA_D1 | 10.42 ± 3.53; 11.5 (8.25; 12.75) | 8.33 ± 2.72 (3 NA); 8 (7; 10.5) | W = 188<br>p = 0.07023 |

## References

1. Cecconi M, Evans L, Levy M, Rhodes A. Seminar Sepsis and septic shock [Internet]. Lancet 2018;6736(April):1–13.

2. Van Vught LAV et al. Incidence, risk factors, and attributable mortality of secondary infections in the intensive care unit after admission for sepsis. JAMA – J. Am. Med. Assoc. 2016;315(14):1469–1479.

3. Soares MP, Teixeira L, Moita LF. Disease tolerance and immunity in host protection against infection [Internet]. Nat. Rev. Immunol. 2017;17(2):83–96.

4. Tsai VWW, Husaini Y, Sainsbury A, Brown DA, Breit SN. The MIC-1/GDF15-GFRAL Pathway in Energy Homeostasis: Implications for Obesity, Cachexia, and Other Associated Diseases [Internet]. Cell Metab. 2018;28(3):353–368.

5. Chung HK et al. Growth differentiation factor 15 is a myomitokine governing systemic energy homeostasis. J. Cell Biol. 2017;216(1):149–165.

6. Hsu J-Y et al. Non-homeostatic body weight regulation through a brainstem-restricted receptor for GDF15 [Internet]. Nature [published online ahead of print: 2017]; doi: 10.1038/nature24042

7. Emmerson PJ et al. The metabolic effects of GDF15 are mediated by the orphan receptor GFRAL [Internet]. Nat. Med. 2017;(August):1–9.

8. Mullican SE et al. GFRAL is the receptor for GDF15 and the ligand promotes weight loss in mice and nonhuman primates [Internet]. Nat.Med. [published online ahead of print: 2017];(August). doi:10.1038/nm.4392

9. Yang L et al. GFRAL is the receptor for GDF15 and is required for the anti-obesity effects of the ligand [Internet]. Nat.Med. [published online ahead of print: 2017];(August). doi:10.1038/nm.4394

10. Mullican SE, Rangwala SM. Uniting GDF15 and GFRAL: Therapeutic Opportunities in Obesity and Beyond [Internet]. Trends Endocrinol. Metab. 2018; xx:1–11.

11. Maurin A-C et al. GDF15 Provides an Endocrine Signal of Nutritional Stress in Mice and Humans [Internet]. Cell Metab. 2019;29(3):707–718.e8.

12. Johnen H et al. Tumor-induced anorexia and weight loss are mediated by the TGF-β superfamily cytokine MIC-1 [Internet]. Nat. Med. 2007; 13(11):1333–1340.

13. Gordon BS, Kelleher AR, Kimball SR. Regulation of muscle protein synthesis and the effects of catabolic states [Internet]. Int. J. Biochem. Cell Biol. 2013;45(10):2147–2157.

14. Buendgens L, Yagmur E, Bruensing J, Herbers U, Baeck C. Growth Differentiation Factor-15 (GDF-15) is a predictor of mortality in critically ill patients with sepsis. Dis. Markers 2017;15.

15. Soares MP, Teixeira L, Moita LF. Disease tolerance and immunity in host protection against infection. Nat. Rev. Immunol. 2017;17(2). doi:10.1038/nri.2016.136

16. Kempf T et al. GDF-15 is an inhibitor of leukocyte integrin activation required for survival after myocardial infarction in mice [Internet]. Nat. Med. 2011;17(5):581–588.

17. Chung HK et al. GDF15 deficiency exacerbates chronic alcohol- and carbon tetrachloride-induced liver injury [Internet]. Sci. Rep. 2017;7(1): 1–13.

18. Alves-Filho JC, Spiller F, Cunha FQ. Neutrophil paralysis in sepsis. Shock 2010;34(SUPPL. 1):15–21.

19. Hotchkiss RS, Monneret G, Payen D. Sepsis-induced immunosuppression: from cellular dysfunctions to immunotherapy. [Internet]. Nat. Rev. Immunol. 2013;13(12):862–74.

20. J. A et al. Extracorporeal cell therapy of septic shock patients with donor granulocytes: A pilot study [Internet]. Crit. Care 2011;15(2). doi:10.1186/cc10076

21. Chavey C et al. CXC Ligand 5 Is an Adipose-Tissue Derived Factor that Links Obesity to Insulin Resistance. Cell Metab. 2009;9(4):339–349.

22. Walz A, Strieter RM, Schnyder S, Bern C-, Arbor A. Neutrophil-activating peptide ena- 78(9):129–130.

23. Rittirsch D, Huber-Lang MS, Flierl MA, Ward PA. Immunodesign of experimental sepsis by cecal ligation and puncture [Internet]. Nat Protoc 2009;4(1):31–36.

24. Kolmogorova D, Murray E, Ismail N. Monitoring Pathogen-Induced Sickness in Mice and Rats. Curr. Protoc. Mouse Biol. 2017;7(2):65–76.

